# Q-STARZ: Quantitative Spatial and Temporal Assessment of Regulatory element activity in Zebrafish

**DOI:** 10.1101/2020.09.13.290460

**Authors:** Shipra Bhatia, Dirk Jan Kleinjan, Kirsty Uttley, Anita Mann, Nefeli Dellepiane, Wendy A. Bickmore

## Abstract

Noncoding regions of the genome harbouring cis-regulatory elements (CREs) or enhancers drive spatial and temporal gene expression. Mutations or single nucleotide polymorphisms (SNPs) in enhancers have been widely implicated in human diseases and disease-predispositions. However, our ability to assay the regulatory potential of genetic variants in enhancers is currently very limited, in part because of the need to assay these elements in an appropriate biological context. Here, we describe a method for simultaneous quantitative assessment of the spatial and temporal activity of wild-type (Wt) and disease-associated, mutant (Mut) human CRE alleles using live imaging in zebrafish embryonic development. We generated transgenic lines harbouring a dual-CRE dual-reporter cassette in a pre-defined neutral docking site in the zebrafish genome. Using this single transgenic cassette, the functional activity of each CRE allele is reported via expression of a specific fluorescent reporter, allowing the simultaneous visualisation of the activity of both alleles. This can reveal where and when in embryonic development the wild-type allele is active and how this activity is altered by the disease-associated mutation.

## INTRODUCTION

Mutations or single-nucleotide polymorphisms (SNPs) in noncoding regions of the human genome functioning as cis-regulatory elements (CREs) or enhancers have been widely implicated in human diseases and disease-predispositions [1, 2]. Disease-associated sequence variation in enhancers can alter the transcription factor (TF) binding sites, leading to aberrant enhancer function and altered target gene expression [2]. Application of next generation sequencing technologies combined with molecular genetic approaches has enabled widespread identification of presumptive CREs and associated putative pathogenic mutations in patient cohorts [3]. However, incomplete understanding of the TF binding potential of CREs impedes the functional assessment of pathogenicity of these variants compared to genetic variants in coding regions. Thus, determining how mutations in the vast stretches of the human noncoding genome contribute to disease and disease-predisposition remains a huge unmet challenge.

Functional analysis of enhancer activity, and assessment of the impact of disease-associated variation on this activity, depends on the availability of the right TFs in the right stoichiometric concentrations, which is only precisely captured *in vivo*. Enhancer-reporter transgenic assays have been widely employed in a variety of model organisms, including the mouse, for *in vivo* assessment of enhancer function [4–8].These assays however can be affected by the random integration of transgenes and have limited application for studying the temporal aspects of enhancer function over the complete time course of embryonic development since, for example, live imaging is challenging due to the opaqueness of the mouse embryo and its *in utero* embryonic development.

Zebrafish (*Danio rerio*) is a highly suitable *in vivo* vertebrate model for visualizing tissue-specific enhancer activity. Robust transgenesis methods allow rapid generation of transgenic lines yielding transparent embryos which develop externally [9, 10]. The activities of a large number of putative human and mouse CREs have been assessed in transgenic zebrafish models, irrespective of the primary sequence conservation of the mammalian CREs in zebrafish [5, 11–16]. However these assays were based on Tol2-recombination which mediates random integration of the CRE-reporter cassette in the zebrafish genome [17]. The measured CRE activities were strongly influenced by the variable site and copy number of integrations, necessitating analysis of each element in multiple transgenic lines and precluding quantitative assessment of CRE activities. These biases can be alleviated by targeted integration of the transgenic cassette into pre-defined neutral sites in the zebrafish genome using phiC31-mediated recombination [18–20].

Previously, we developed a system in which dual fluorescence CRE-reporter zebrafish transgenics allow for direct comparison of the *in vivo* spatial and temporal activity of wildtype (Wt) and putative SNP/mutation (Mut) bearing CREs in the same developing embryo [5]. The functional output from each CRE version (Wt/Mut) is visualized simultaneously as eGFP or mCherry signal within a live developing embryo bearing both transgenes. This enables unambiguous comparison of the activity of both wildtype and mutant CREs in a developmental context, simultaneous assessment of multiple separate elements for subtle differences in spatio-temporal overlap, and the validation of putative TFs by analysing the effect of morpholino-mediated depletion of the putative TF on CRE activity [5]. The assay had clear advantages over other conventional CRE-reporter transgenic assays, notably rapid, unambiguous detection of subtle differences in CRE activities using a very low number of animals. However, as the CRE alleles were on separate constructs randomly integrated into the zebrafish genome, the assay was not suitable for quantitative assessment of altered CRE activity. Furthermore, multiple transgenic lines had to be analysed for each CRE to eliminate any bias arising from the site of integration.

Here we describe Q-STARZ (**Q**uantitative **S**patial and **T**emporal **A**ssessment of **R**egulatory element activity in **Z**ebrafish), a new and significantly improved design of our previous transgenic reporter assay, based upon targeted integration of a dual-CRE dual-reporter cassette into a pre-defined site in the zebrafish genome (Fig. 1). A unique feature of this design is the single transgenic cassette containing both wildtype and mutant CREs, separated by strong insulator sequences, with the transcriptional potential of both CREs read out as expression of different fluorescent proteins. Qualitative and quantitative activity of the two CRE alleles is analyzed from eGFP/mCherry fluorescence in real-time by live imaging of embryos obtained from the founder (F0) lines bearing the dual CRE dual-reporter cassette. This allows robust, unbiased assessment of spatial and temporal activities of both CREs using a single transgenic line, thereby reducing animal usage by up to 75% compared to the previous design. We utilize disease-associated mutations in well characterized CREs from the *PAX6* and *SHH* loci to demonstrate the salient features of the Q-STARZ method.

## RESULTS

### Targeted integration of dual-CRE dual-reporter transgenic cassette in the zebrafish genome

Analysis of enhancer activities in conventional reporter assays in zebrafish suffer from bias arising from position effects due to the random integration of the transgene at Tol2 sites naturally distributed at a low frequency throughout the zebrafish genome [9, 17]. Most assay designs also harbour only one CRE per transgene introducing ambiguity in the analysis when comparing CREs with highly similar activities or subtle changes in sequence (e.g. disease-associated mutations or SNPs). Q-STARZ is a versatile, robust and cost-effective analysis pipeline designed to alleviate both these limitations (Fig. 1).

We first generated ‘landing lines’ harbouring phiC31 attB integration sites at inert positions in the zebrafish genome. Using Tol2-mediated transgenesis, we integrated ‘landing pads’ at random sites in the zebrafish genome (Fig. 1, supplementary fig. 1). To visualise successful integration events, the landing pads contain ‘tracking CREs’ (Supplementary table 1) driving expression of a ‘tracking reporter gene’. These CREs had previously well-characterised activities, enabling us to select transgenic lines devoid of bias arising from the site of integration [5]. We assessed reporter gene expression in F1 embryos derived from several independent F0 transgenic lines for each tracking CRE (Fig. 2, supplementary table 2). F1 embryos in which the activity of CRE was not influenced by the site of integration were raised to adulthood to establish ‘landing lines’ presumed to be harbouring the phiC31 attB sites in an inert position of the zebrafish genome (Fig. 1B). The CRE activities were observed to be highly influenced by the site of integration in F1 embryos derived from founder lines bearing *SOX9*-CNEa and *Pax6*-SIMO CREs, but we obtained landing lines with clean, single-site integration using *Shh*-SBE2 as the tracking CRE (Fig. 2, supplementary table 2). *Shh*-SBE2 is a forebrain enhancer driving *Shh* expression in the hypothalamus [21]. Based on these observations, we decided to use the *Shh*-SBE2 landing line for all subsequent experiments described in this study. The precise integration site of the landing pad in the *Shh*-SBE2 line was determined using ligation mediated PCR (LM-PCR) (Supplementary fig. 2).

In the second part of the Q-STARZ pipeline, we generated a ‘dual-CRE dual-reporter replacement construct’ containing two CRE-reporter cassettes separated from each other by strong insulator sequences (Fig. 1A, supplementary fig 1B). The replacement construct was co-injected with mRNA encoding phiC31 integrase into F2 embryos derived from the *Shh*-SBE2 landing line. Recombination-mediated cassette exchange between the attB sites on the landing pad construct and attP sites on the replacement construct integrates a single copy of the dual-CRE dual-reporter cassette at the predefined site in the zebrafish genome (Fig. 1B). Injected embryos were scored for loss of *Shh*-SBE2 driven CRE activity in the forebrain and gain of mosaic eGFP and mCherry signals from the replacement CRE-reporter cassette. Selected embryos were raised to sexual maturity to establish independent founder transgenic lines for each replacement cassette analysed in this study (Supplementary table 2). Activity of both CREs in the replacement construct was visualised simultaneously as eGFP or mCherry signals by live imaging of F1 embryos derived from outbreeding the founder lines with wild-type zebrafish (Fig. 1B). Detailed protocols for the various steps described in this section are provided in materials and methods.

### Robust, quantitative assessment of CRE activity using Q-STARZ

A key feature of Q-STARZ is the simultaneous assessment of activities of the two CREs present on the replacement cassette. In order to prevent cross-talk between the two enhancers, a well-characterised insulator sequence from the chicken genome, chicken β-globin 5’HS4 (cHS4) [22, 23] was placed between the two CRE-reporter cassettes (Fig. 1A, supplementary fig. 1). We optimised the assay using replacement constructs bearing two CREs with previously well-characterised tissue-specific activities from the *PAX6* regulatory domain (Fig.3). PAX6 is a transcription factor with vital pleiotropic roles in embryonic development [4, 24, 25]. Over 30 CREs have been characterised upstream and downstream of the *PAX6* transcription start site which coordinate precise spatial and temporal *PAX6* expression in the developing eyes, brain and pancreas [2]. We selected *PAX6*-7CE3 and *PAX6*-SIMO for this analysis as they have well established and highly distinct tissue-specific activities during zebrafish embryogenesis. *PAX6*-7CE3 drives expression in the hindbrain and neural tube from 24hpf-120hpf, while *PAX6*-SIMO activity is in developing lens and forebrain from 48hpf-120hpf [14, 16] (Supplementary table 1).

When the two CRE-reporter cassettes were separated by a ‘neutral’ sequence – a randomly selected region from the mouse genome with no insulator activity, we observed complete crosstalk of the two CRE activities (Fig. 3A, supplementary fig. 3, and supplementary table 2). We also performed a dye-swap experiment wherein the eGFP and mCherry reporters were swapped between the two CREs. We observed no significant difference in CRE activities in the dye swap experiment, indicating no bias was introduced in the analysis by varying signal intensities from the two fluorophores used (Supplementary fig. 3). Next, we substituted the neutral sequence with one, two or three tandem copies of the cHS4 insulator. Enhancer blocking activity of this insulator has been attributed to its ability to bind CTCF [26]. Crosstalk between the two enhancer-reporter cassettes was progressively reduced with increasing copies of cHS4, with complete insulation achieved in replacement cassettes bearing three copies (3xcHS4) (Fig. 3A, supplementary fig. 4–6). We quantified the effects of the presence of insulator sequences by measuring eGFP and mCherry intensities in the expressing tissues at all stages of embryonic development in multiple embryos for each of the construct analysed. The quantification analysis was focussed on lens and hindbrain tissues as we observed expression in these sites consistently in all the lines analysed (Supplementary table 2). This analysis confirmed that as the number of copies of the insulator increases there is progressively restricted expression of the reporters towards expression only in the activity domains of their associated CRE (Fig. 3B).

### Dissecting spatial and temporal dynamics of CREs with highly overlapping activities using live imaging

A salient feature of Q-STARZ is the ability to simultaneously visualise the activity of both CREs on the replacement cassette in the same developing zebrafish embryo in real-time using live imaging. To establish proof of principle, we investigated the precise spatial and temporal activities of two CREs from the *Shh* locus, *Shh*-SBE2 and *Shh*-SBE4, previously demonstrated to have highly similar domains of activity in the developing forebrain of mouse embryos (Fig. 4A) [21, 27]. Our analyses revealed unique, as well as overlapping, domains of activity of both CREs in the early stages of forebrain development (~24hpf-50hpf) (Fig. 4B, supplementary movie 1). However, from ~60hpf-120hpf, the activities of both CREs are observed in completely distinct domains of the developing forebrain with no overlapping activity observed. *Shh*-SBE2 was active in the rostral part of forebrain while *Shh*-SBE4 activity was restricted to caudal forebrain (Fig. 4B, supplementary movie 2). This analysis highlights the importance of simultaneous visualization of CRE activities in the developing embryo to define the precise spatial and temporal activity of each CRE.

### Robust assessment of the effects of disease-associated mutations on CRE activity

As well as qualitative comparison of activity between two different CREs, a key strength of the Q-STARZ pipeline is its suitability for discerning the precise effects of disease-associated mutations or SNPs within a specific CRE. We tested this in SBE2, a regulatory element that controls *SHH* expression in the developing forebrain, using a point mutation (C>T) identified in a patient with holoprosencephaly, and shown to abrogate the activity of SBE2 in the rostral hypothalamus of the mouse (Fig. 5A) [5, 27]. We simultaneously visualised the activities of the human Wt(C) and Mut(T) SBE2 alleles in our dual-CRE dual-reporter system by live imaging of transgenic zebrafish embryos from 24hpf – 72hpf (Fig. 5B, supplementary movie 3). We detected no difference in the activities of the two alleles in very early development until ~40hpf. However, from ~ 48-72hpf, activity of the alleles started to diverge. Expression driven by the Wt allele was observed in the developing rostral and caudal hypothalamus of transgenic embryos while the Mut allele was only active in the caudal hypothalamus, indicating that the mutation disrupts rostral activity of the SBE2. Upon quantification of reporter gene expression associated with each allele, we observed no significant difference in activity was observed between the two alleles at 28hpf. However at later stages of development (48hpf and 72hpf) the mutant allele failed to drive reporter gene expression in the rostral hypothalamus, and had significantly weaker activity in the caudal hypothalamus (Fig. 5C).Our analysis thus unambiguously and precisely uncovered where and when in embryonic development the mutation associated with holoprosencephaly alters the enhancer activity of SBE2.

## DISCUSSION

The noncoding region of the human genome is estimated to contain approximately one million enhancers [28, 29]. The widespread application of whole-genome sequencing for understanding genetic diseases (rare, common and acquired – i.e. cancer), combined with genome-wide identification of chromatin signatures associated with active enhancers, has led to the identification of a large number of putative enhancers with disease-associated or disease risk-associated sequence variation [1, 2, 30, 31]. A complete understanding of how these sequence changes alter enhancer function is a necessary first step towards establishing roles of the CREs in the aetiology of the associated disease. Thus there is a pressing need for rapid, cost effective assays for robust unambiguous comparisons of mutant CRE alleles with the activities of wild-type alleles. Importantly, this has to be done in the appropriate context, relevant to the biology of the associated disease. CRE activity depends on precise stoichiometric concentrations of specific transcription factors, which is only achieved in the right physiological context inside a developing embryo or in cell lines that closely model the cellular phenotypes of developing tissues [32, 33].

The Q-STARZ assay we describe here is highly versatile and enables unambiguous assessment of human tissue-specific CRE function *in vivo* at all stages of early embryonic development in a vertebrate model system. A distinctive feature of the assay is the targeted integration of a single transgenic cassette bearing two independent CRE-reporter units into a pre-defined inert site in the zebrafish genome. We establish an analysis pipeline that enables simultaneous robust qualitative and quantitative analysis of enhancer function without any bias from position effects or copy number variation between the two CREs analysed.

Similar methods of targeted integration of enhancer-reporter transgenic cassettes have been developed for zebrafish as well as mouse models [18, 34]. Q-STARZ however offers a unique advantage when analysing the effects of disease-associated sequence variation on CRE function by enabling direct comparisons of activities of wild-type and mutant alleles inside the same, transparent developing embryo using live imaging.

Using CREs with previously well-established activities we demonstrate that inclusion of three tandem copies of a strong insulator sequence in the replacement construct robustly prevents cross talk between the two CREs analysed. This feature enables direct comparisons of the spatial and temporal dynamics of both CREs by simultaneous visualization of functional outputs in live embryos at all stages of development. We demonstrate here that this can uncover the precise sites and time points in embryonic development where the CRE functions are unique and where they overlap with each other. Q-STARZ is therefore an ideal tool for generating a detailed cell-type specific view of CRE usage during embryonic development. This will enhance understanding of the roles of CREs in target gene regulation, particularly for the complex regulatory landscapes of genes with key roles in development like *PAX6* and *SHH*. Analysis of CREs derived from these loci in conventional transgenic assays has revealed multiple CREs apparently driving target gene expression in the same or highly overlapping tissues and cell-types. This has led to the concept of redundancy in enhancer function conferring robustness of expression upon genes with key roles in embryonic development [35–37]. However, our analysis of the *Shh*-SBE2 and SBE4 enhancers, previously reported as forebrain enhancers with overlapping functions, reveals subtly distinct spatial and temporal activity domains of each enhancer during development. Based on these results, we hypothesize that there are small but important differences in the timing of action or precise localisation in cell-types within the forebrain where these enhancers exert their roles that are overlooked when analysed independently in conventional transgenic assays.

Finally, we demonstrate that Q-STARZ can robustly detect differences in activities of mutant and wild-type CRE alleles. Live imaging of transgenic embryos carrying a reporter cassette with a previously validated disease-associated point mutation in the *SHH*-SBE2 enhancer revealed the loss-of-activity of the mutant allele in the rostral hypothalamus compared to the wild-type CRE. This recapitulates a similar loss of rostral activity of the SBE2 mutant that has been previously reported in mouse transgenic assays [27]. However, since we could visualise the activities of both the wild-type and mutant alleles simultaneously in the same embryos in real-time, we were able to determine the precise time point in development when the mutation affects CRE function. We propose that Q-STARZ will be a powerful tool to define the precise cell-types and stages of development where CRE function is affected by mutations or SNPs identified by GWAS and other studies, thus this could significantly improve our ability to discern potentially pathogenic and functional sequence variation from background human genetic variation, which is currently a major challenge for human genetics.

## MATERIALS AND METHODS

### Ethics statement

All zebrafish experiments were approved by the University of Edinburgh ethical committee and performed under UK Home Office license number PIL PA3527EC3; PPL IFC719EAD.

### Generation of landing pad and dual-CRE dual-reporter replacement vectors

All the constructs in this study were generated using the Gateway recombination cloning system (Invitrogen). PCR primers with suitable recombination sites were used for amplification of CREs from the genomic DNA (Supplementary table 1). The PCR amplification was performed using Phusion high fidelity polymerase (NEB) and the amplified fragments were cloned in Gateway pDONR entry vectors (pP4P1r or pP2rP3) and sequenced using M13 forward and reverse primers for verification. The recombination sites attached in primers, entry vector for cloning and genomic DNA used in amplification for each CRE are indicated in supplementary table 1. For generating the landing pad vector, pP4P1r entry vector with the tracking CRE and pDONR221 entry vector containing a gata2-eGFP [5] were recombined with a destination vector with a Gateway R4-R2 cassette flanked by phiC31_attB1/B2 and Tol2 recombination sites (Supplementary fig. 1A). The details of the tracking CREs are provided in supplementary table 1. The replacement vector was generated via three-way gateway reaction as described in supplementary fig. 1B. The test CREs were cloned either in pP4P1r or pP2rP3 entry vectors and the insulator sequences and neutral sequence was cloned in pDONR221. For generating constructs with multiple copies of the insulator sequence, the sequences were first cloned in tandem in TOPO TA Cloning Kit (Thermo Fischer Scientific, cat no 451641). Plasmids containing one, two or three copies of the insulator sequence were used as templates for amplification of products suitable for cloning in pDONR221. The destination vector was synthesized by Geneart and contained a Gateway R4-R3 cassette flanked by phiC31_attP1/P2 recombination sites and minimal promoter-reporter gene units (gata2-eGFP and gata2-mCherry). Details of each construct generated in the manuscript are provided in supplementary table 1 and complete vector maps for all the constructs would be available on request.

### Generation of zebrafish transgenic lines

Zebrafish were maintained in a recirculating water system according to standard protocols [38]. Embryos were obtained by breeding adult fish of standard stains (AB) and raised at 28.5°C as described [38]. Embryos were staged by hours post fertilization (hpf) as described [39]. Final CRE-reporter plasmids were isolated using Qiagen miniprep columns and were further purified on a Qiagen PCR purification column (Qiagen), and diluted to 50 ng/ml with nuclease free water. Tol2 transposase mRNA and phiC31 integrase mRNA were synthesized from a NotI-linearized pCS2-TP or pcDNA3.1 phiC31 plasmid, respectively [40, 41] using the SP6 mMessage mMachine kit (Ambion), and final RNA diluted to 50 ng/ml. Equal volumes of the reporter construct(s) and the transposase RNA were mixed immediately prior to injections. 1–2 nl of the solution was micro-injected per embryo and up to 200 embryos were injected at the 1- to 2-cell stage. Embryos were screened for mosaic fluorescence at 1–5 days post-fertilization i.e. 24–120 hpf and raised to adulthood. Germline transmission was identified by outcrossing sexually mature F0 transgenics with wild-type fish and examining their progeny for reporter gene expression/ fluorescence. F1 embryos from 3–5 F0 lines showing the best representative expression pattern for each construct were selected for confocal imaging (Fig. 1B). A few positive embryos were also raised to adulthood and F1 lines were maintained by outcrossing. A summary of the number of independent lines analysed for each construct and their expression sites is included in supplementary table 2.

### Mapping of transgene integration site in the landing lines

Transgenic embryos obtained from lines harbouring the landing pad vectors were sorted into eGFP-positive and eGFP-negative groups. Genomic DNA was purified from ~ 100 pooled embryos using QIAGEN DNeasy blood and tissue kit (Cat No./ID: 69504). Ligation-mediated PCR (LM-PCR) [42] was used for mapping the landing pad integration site using previously published protocol [43]. 1μg of genomic DNA was digested with either NlaIII, BfaI or DpnII and purified using a Qiagen QIAquick PCR purification kit (Cat No./ID: 28104). A 5μl aliquot was added to a ligation reaction containing 150 μmoles of a double stranded linker. Ligations were performed using high concentration T4 ligase (NEB, M020S) at room temperature for 2-3 hours. The first round of the nested PCR was performed using linker primer 1 with either Tol2 Left 1.1 or Tol2 Right 1.1, using the following cycling conditions: 94 °C (15 s)–51 °C (30 s)–68 °C (1 min), 25–30 cycles. Second round nested PCR was then performed using linker primer 2 with either Tol2 Left 2.1 or Tol2 Right 2.1 and the following cycling conditions: 94 °C (15 s)–57.5 °C (30 s)–68 °C (1 min), 25– 30 cycles. The PCR products were resolved by electrophoresis on a 3% agarose gel and the products selectively amplified in samples derived from eGFP-positive embryos were cloned and sequenced. Sequences flanking the Tol2 arms were used to search the Ensembl *Danio rerio* genomic sequence database to position and orient the insert within the zebrafish genome. The sequences of the linker oligos and primers used are provided in supplementary table 1.

### Imaging of zebrafish transgenic lines

Embryos for imaging were treated with 0.003% PTU (1-phenyl2-thio-urea) from 24hpf to prevent pigmentation. Embryos selected for imaging were anaesthetized with tricaine (20-30mg/L) and mounted in 1% low-melting point (LMP) agarose. Images were taken on a Nikon A1R confocal microscope and processed using A1R analysis software. Time-lapse imaging was performed on an Andor Dragonfly spinning disk confocal, and processed using Imaris (Bitplane, Oxford Instruments) and ImageJ. Embryos mounted in 1% LMP were covered with tricaine solution and held in a chamber at 28.5°C

### Quantification of imaging data

eGFP and mCherry signal intensities were quantified in selected regions of expression in images acquired from F1 transgenic embryos using ImageJ software. Measurements were taken from at least five independent embryos for each line. Mean fluorescence intensity ratios (eGFP/ mCherry, G/C or mCherry/ eGFP, C/G) were computed for each expression domain. Average of mean fluorescence intensity ratios was computed using measurements from independent embryos derived from each line for each expression domain and plotted as shown in fig. 3b and 5c. The level of significance (p-value) of differences in average mean florescence intensity ratios in expressing tissues between different transgenic lines was computed using two-tail student t-test. Raw values of the data plotted are provided in supplementary table 3.

**Fig. 1:**
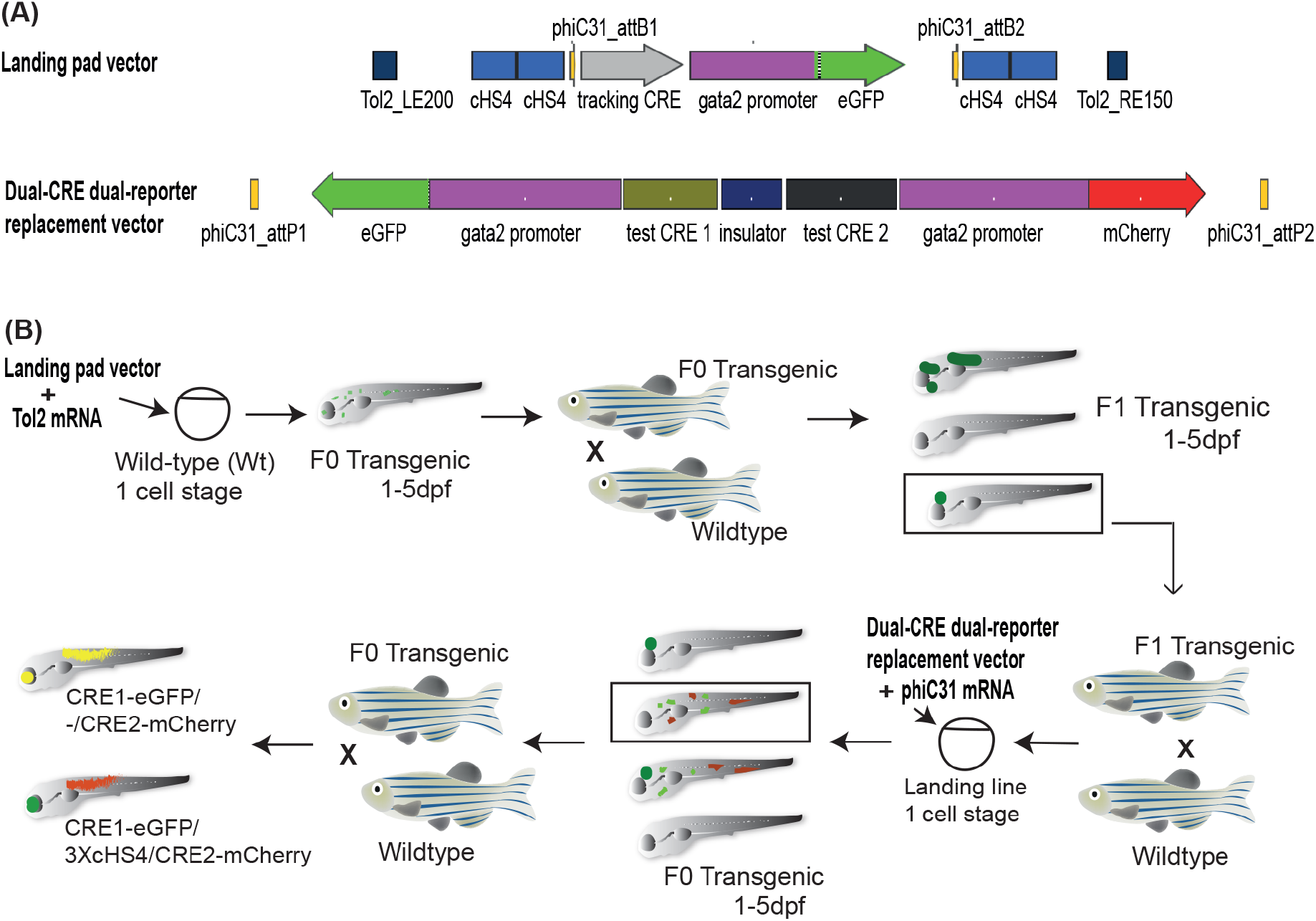
Q-STARZ pipeline for Quantitative Spatial and Temporal Assessment of Regulatory element activity in Zebrafish. (A) Simplified maps of the landing pad and dual-CRE dual-reporter replacement vectors. (B) Diagrammatic representation of the Q-STARZ pipeline. Top panel (left to right); scheme for generating stable transgenic ‘landing lines’. The landing pad vector is co-injected with Tol2 mRNA into one-cell stage wild-type embryos. Tol2-mediated recombination integrates the landing pad containing phiC31-attB sites flanking the tracking CRE-reporter cassette (*SHH*-SBE2, a CRE active in the developing forebrain, driving eGFP depicted here) at random locations in the zebrafish genome. F0 embryos showing mosaic eGFP expression are raised to adulthood. F1 embryos obtained by outcrossing F0 lines with willd-type zebrafish were screened for tracking CRE driven reporter gene expression (*SHH*-SBE2 driven eGFP expression in the developing forebrain). Embryos where eGFP expression was only observed in the expected activity domain of the tracking CRE were selected and raised to adulthood to establish stable ‘landing lines’. Bottom panel (right to left); scheme for replacing the tracking cassette in the landing line with the dual-CRE dual-reporter cassette containing the enhancers to be assessed for spatio-temporal activity. Replacement vector and mRNA coding for phiC31 integrase are injected in one cell stage embryos derived from outcrossing F1 landing line with wild-type fish. Injected embryos were selected for loss of tracking CRE (*SHH*-SBE2) driven eGFP fluorescence in forebrain and mosaic expression of both eGFP and mCherry resulting from the test CREs in replacement cassette. F0 transgenic lines were established from selected embryos and eGFP and mCherry expression was observed by imaging F1embryos derived from outcrossing these lines with wild type fish. Signals from both reporter genes were observed in the activity domains of both CREs in F1 embryos bearing the replacement constructs with ‘neutral’ sequence between the two CRE-reporter units (yellow signal seen in expressing tissues in the merge channel). However, eGFP and mCherry expression were restricted to tissues where the associated CREs are active upon inclusion of three copies of cHS4 (3XcHS4) between the two CRE-reporter units in the replacement construct.

**Fig. 2:**
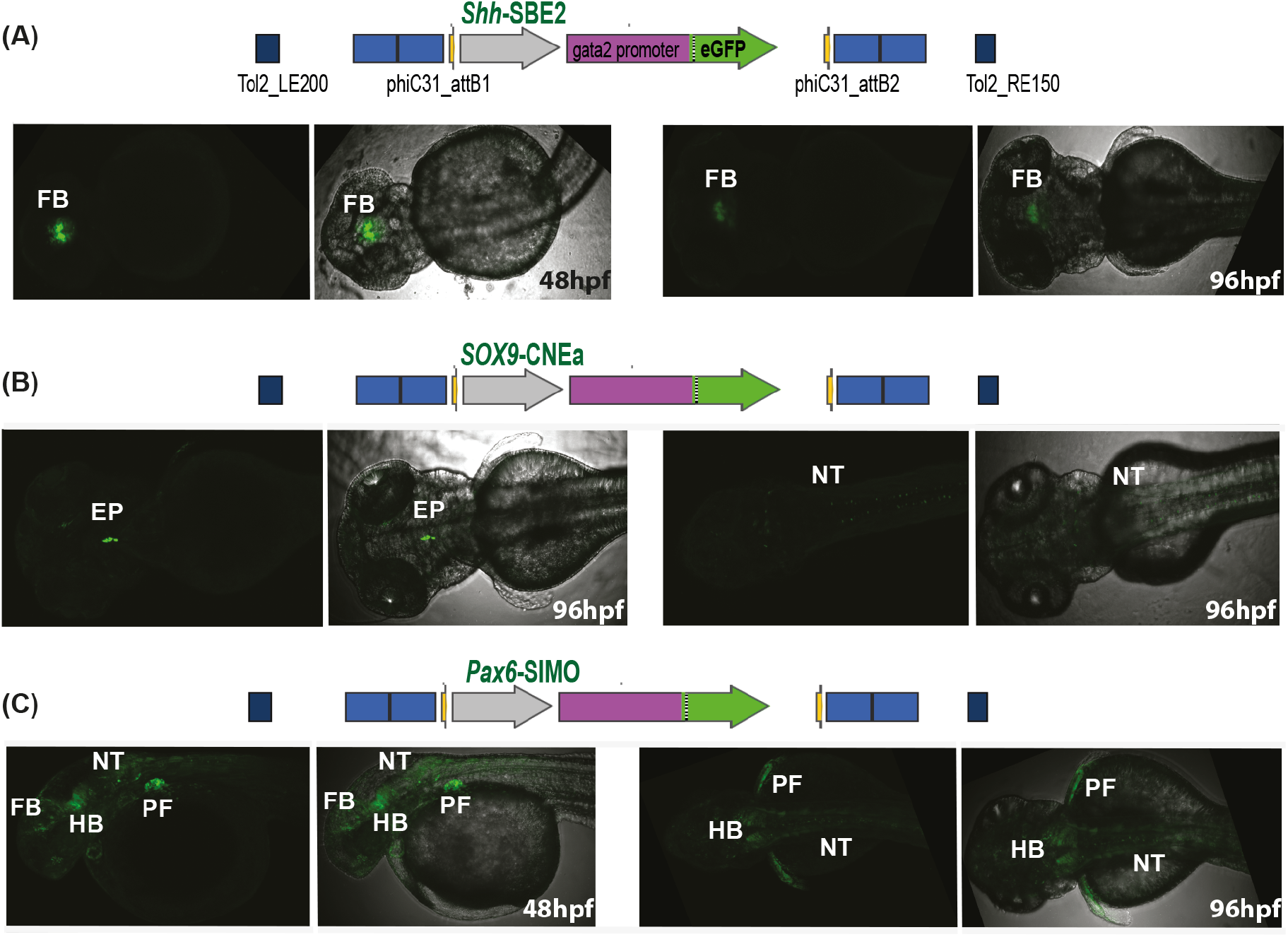
Characterisation of landing lines using tracking CRE driven reporter gene expression. Schematic at the top of each image panel depicts the tracking CRE used in the landing line. (A) CRE activity observed exclusively in the forebrain, the established activity domain of *Shh*-SBE2, in F1 embryos from landing line with *SHH*-SBE2-eGFP tracking cassette. (B-C) Activities of the *SOX9*-CNEa and *Pax6*-SIMO CREs in the landing pad are highly influenced by the site of integration indicated by eGFP expression in multiple tissues. FB-Forebrain; EP-Ethmoid plate; NT-Neural tube; HB-Hindbrain; PF-Pectoral fin; hpf-hours post fertilization

**Fig. 3:**
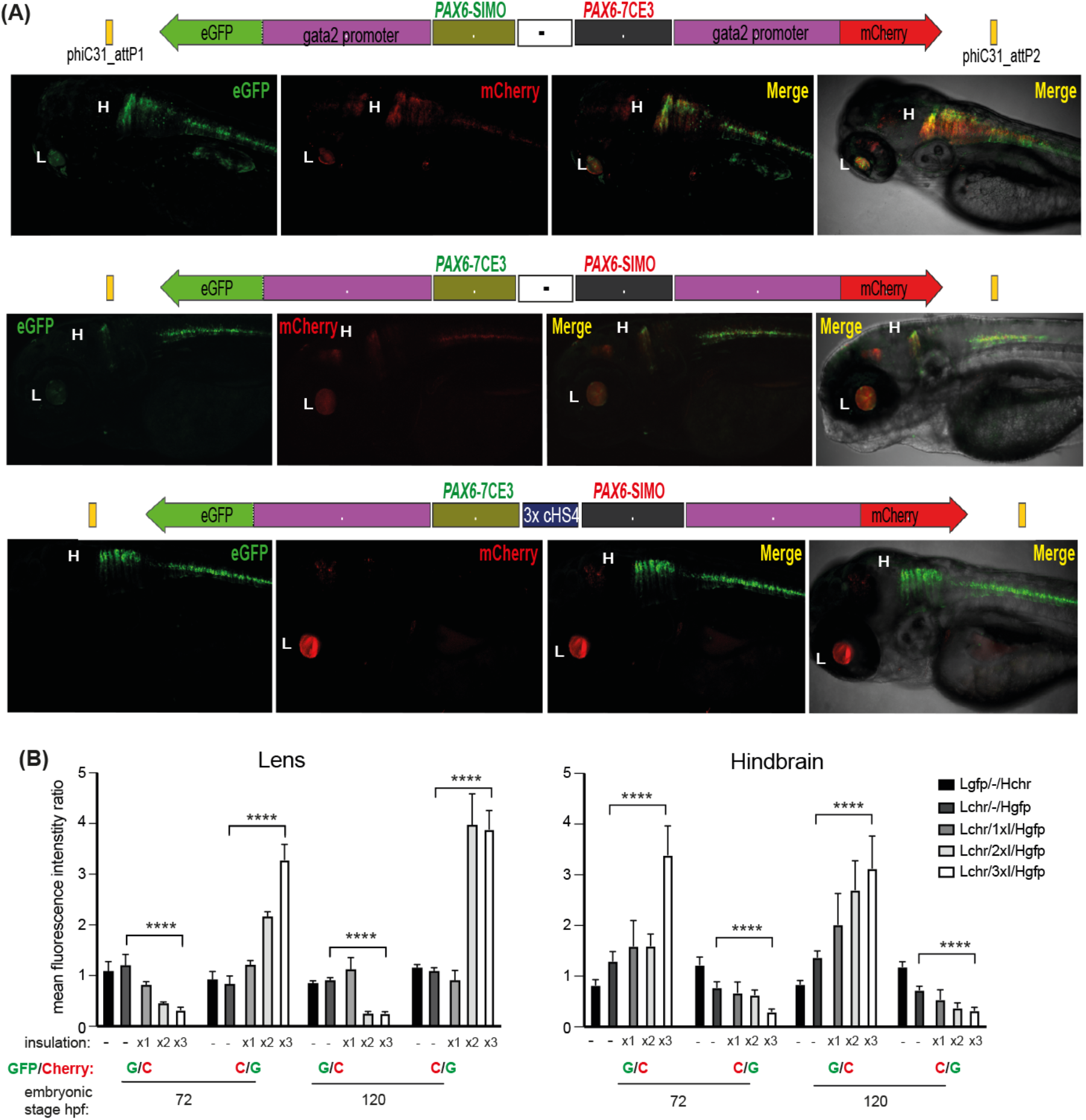
Quantitative assessment of tissue-specific enhancer activity and effect of insulation on crosstalk between CREs in dual-CRE dual-reporter replacement constructs. Replacement constructs carrying well-characterised CREs from the PAX6 locus (*PAX6*-7CE3, hindbrain enhancer and *PAX6*-SIMO, lens enhancer). (A) Confocal images of 96hpf F1 embryos derived from founder lines injected with the replacement cassettes indicated above each image panel. Top two panels: show dye-swap experiment (eGFP and mCherry reporters swapped between the two CREs) with a neutral sequence (-, no insulator activity) between the two CRE-reporter cassettes. eGFP and mCherry expression is observed in both lens and hindbrain indicating complete crosstalk between 7CE3 and SIMO CREs. Bottom panel: Inclusion of three copies of the well-characterised chicken β-globin 5’HS4 (3XcHS4) insulator restricts the activities of each CRE to their respective specific domains (eGFP in H vs mCherry in L). (B) Average of mean fluorescence intensities ratios (G/C: eGFP/mCherry, C/G: mCherry/eGFP) in the lens and hindbrain at 72 and 120hpf in F1 embryos derived from founders bearing the replacement constructs without (-) or with 1x, 2x or 3x insulator sequences. Each bar indicates average of ratios of mean fluorescence intensities from at least five independent images of embryos bearing the replacement construct indicated. A highly significant change in fluorescence intensity ratios was observed between embryos at the same stage of development harbouring replacement constructs either with neutral sequence (-) and those with three copies of the insulator (3xI). This demonstrates that fluorescence is progressively restricted to the tissue where the associated CRE is active as the number of copies of the insulator increase. L: Lens, H: Hindbrain, hpf: hours post fertilization, ****p<0.0001

**Fig. 4:**
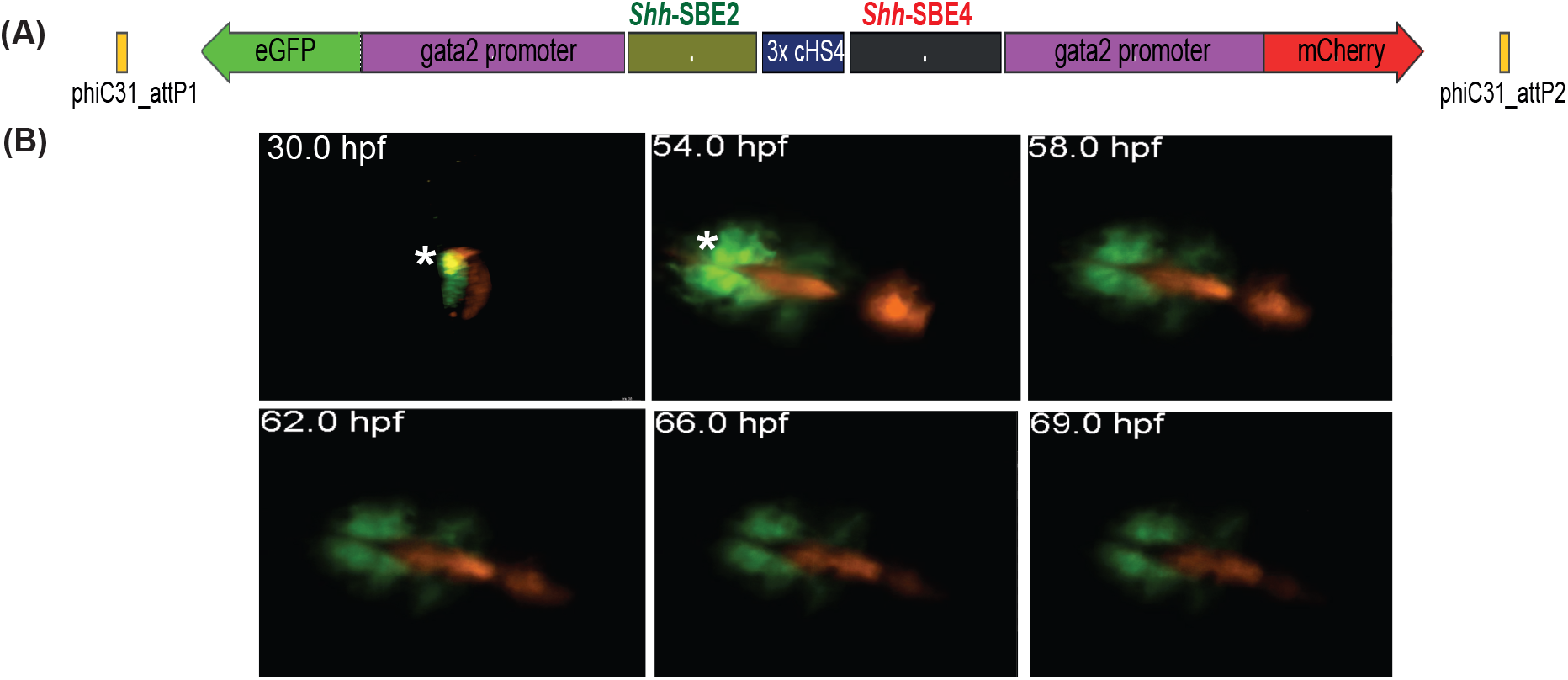
Live imaging of transgenic embryos to detect subtle differences in spatial and temporal activities of two different CREs active in same tissue. (A) Schematic of replacement construct with two enhancers from the mouse *Shh* locus active in developing forebrain (*Shh*-SBE2 and *Shh*-SBE4). (B) Snapshots of live imaging of F1 embryos derived from transgenic lines bearing the replacement construct. Distinct as well as overlapping domains (marked by *) of activities are observed for the two CREs in early stages of development, until about 54hpf. At later stages of embryonic development, the activities of the two forebrain CREs are observed in completely distinct domains.

**Fig. 5:**
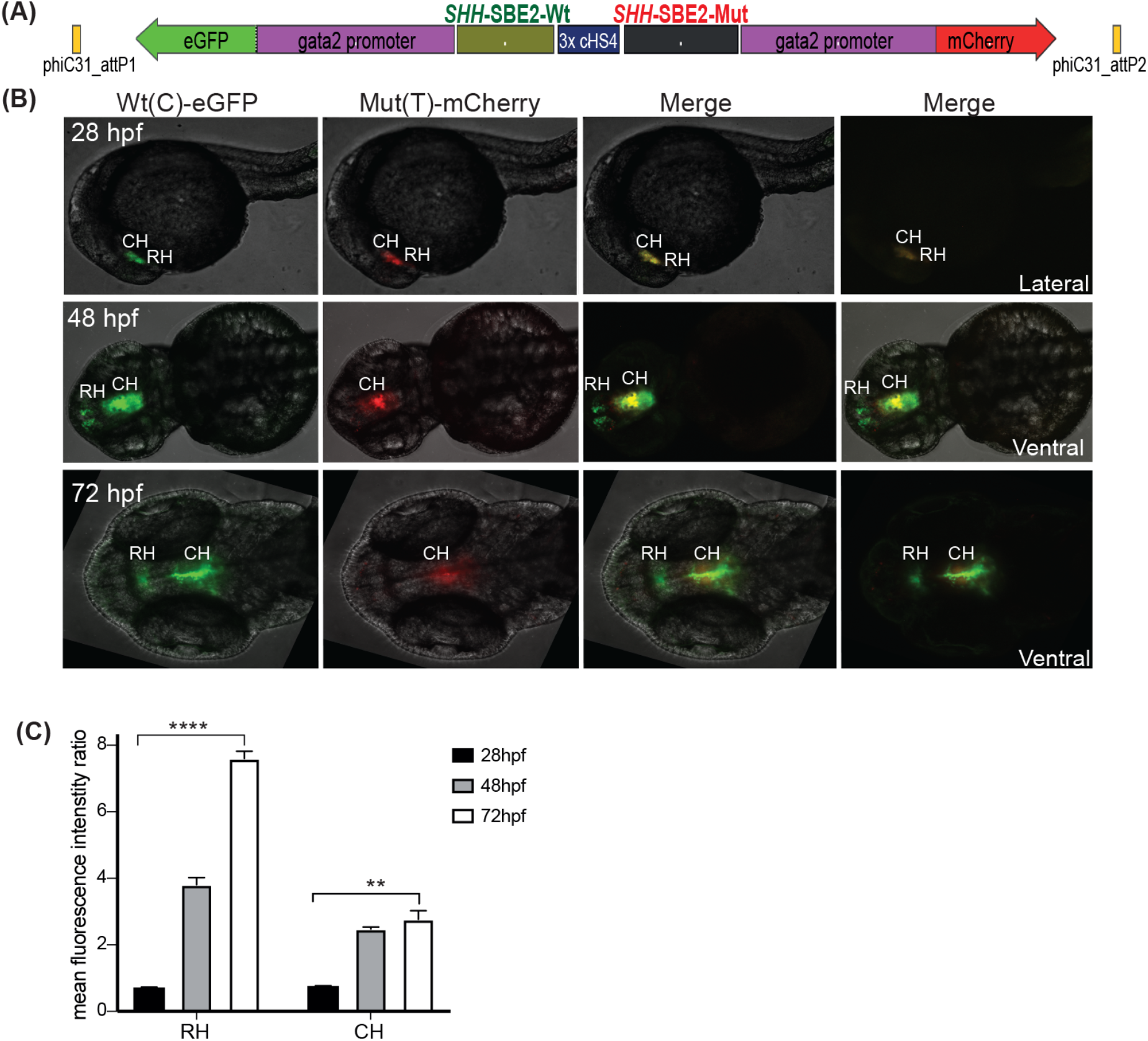
Quantitative assessment of altered CRE activity by disease-associated sequence variation. (A) Replacement construct with *SHH*-SBE2 CRE, wild-type Wt(C) allele driving eGFP and Mut(T) allele bearing a holoprosencephaly associated mutation driving mCherry. (B) Confocal images and (C) histogram showing average of mean fluorescence intensities ratios (eGFP/mCherry) in the rostral (RH) and caudal (CH) hypothalamus for F1 embryos derived from founder lines bearing the replacement construct described in (A). At 28hpf, no significant difference in activity was observed between the two alleles. However, at later stages of development (48hpf and 72hpf) the mutant allele failed to drive reporter gene expression in the RH, and had significantly weaker activity in the CH at 72hpf. ****p<0.0001, **p<0.01

## Acknowledgements

This research was funded by a project grant from Newlife charity for disabled children (632WBI/R43399), a personal fellowship to SB from the Royal society of Edinburgh and Caledonian research fund (632WBI/R45412) and by core funding from the Medical Research Council UK to the Institute of Genetics and Molecular Medicine at the University of Edinburgh.

## Supporting Information

**Supplementary figure 1:**
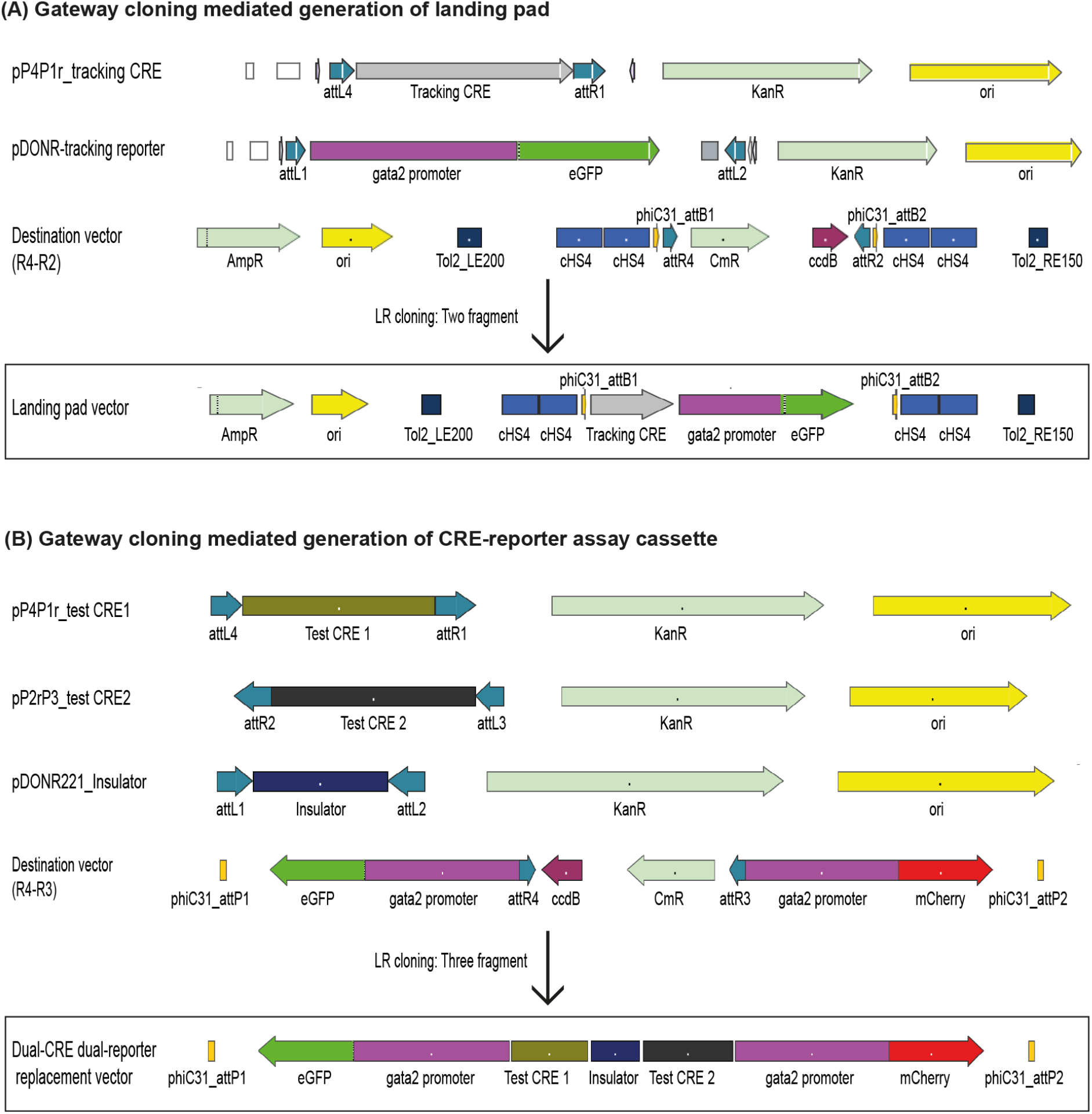
Diagrammatic representation of the gateway cloning strategy used for generating the landing pad and dual-CRE dual-reporter replacement vector. The recombination sites used in each vector and other salient features are indicated on each vector map.

**Supplementary figure 2:**
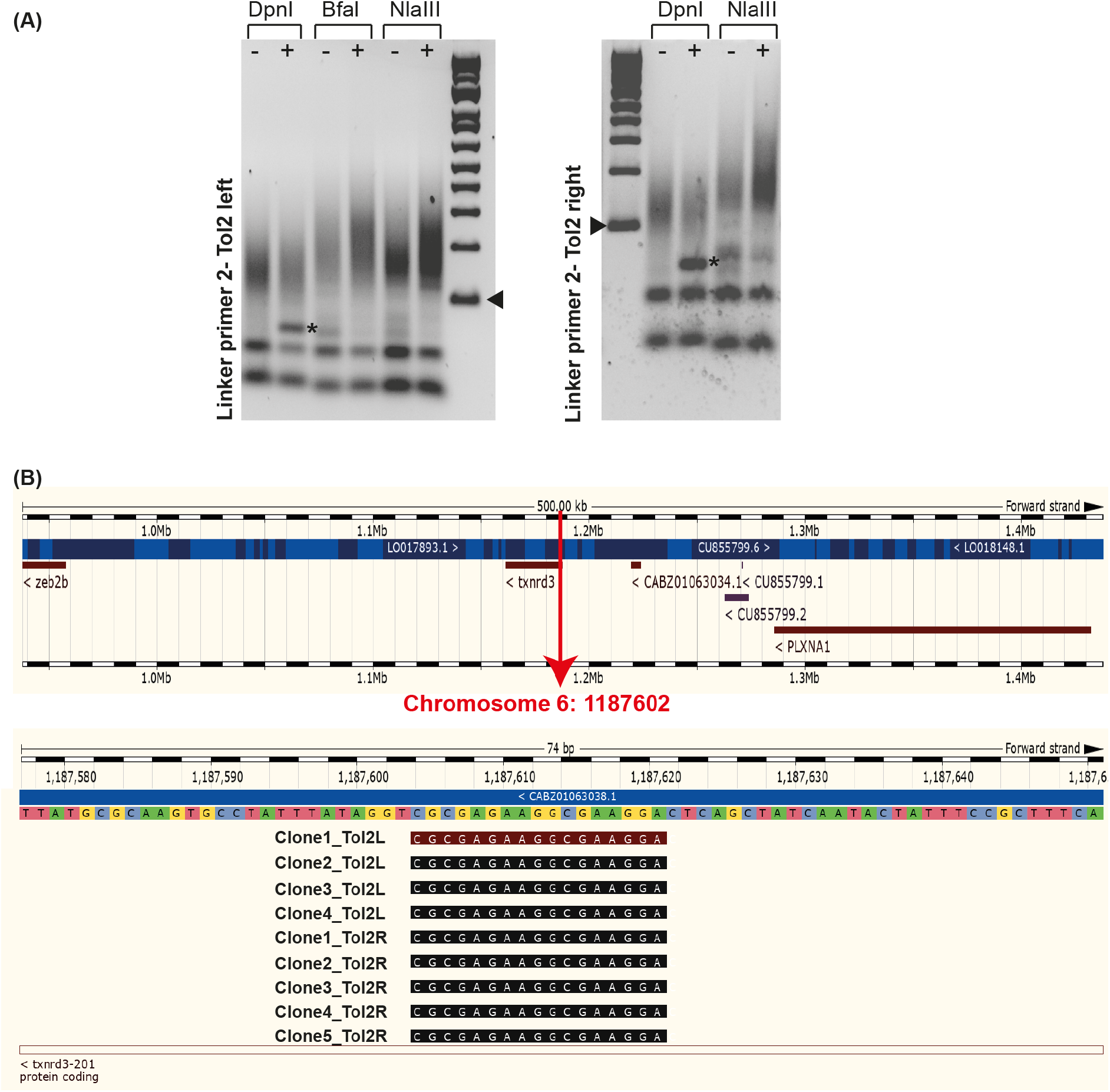
Mapping the site of integration of *Shh*-SBE2 landing line. (A) Unique bands (*) observed in round 2 of PCR amplification of genomic DNA from F1 embryos bearing the landing pad cassette digested with DpnI. (B) Ensembl genome browser snapshot depicting the integration site (red arrow) of the *Shh*-SBE2 landing pad and the sequencing data from the clones bearing the PCR product shown by * in panel A mapped on the integration site.

**Supplementary fig. 3:**
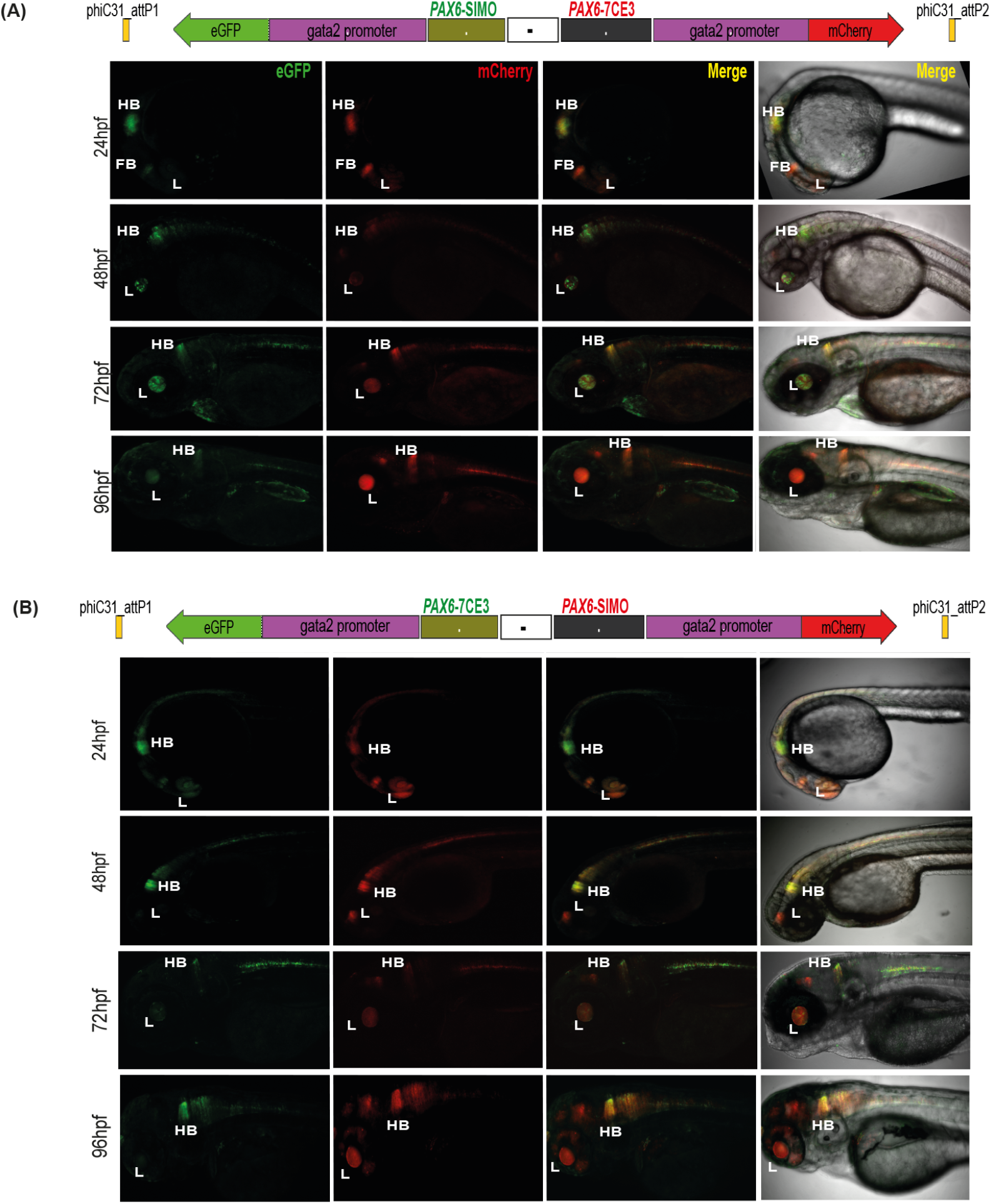
Assessment of tissue-specific CRE activity from dual-CRE dual-reporter replacement constructs with neutral sequence between CREs. Confocal images of F1 embryos (24-96hpf) derived from founder lines injected with the replacement cassettes lacking insulator sequences (-) and containing previously well-characterised CREs from *PAX6* locus (*PAX6*-7CE3, hindbrain enhancer and *PAX6*-SIMO, lens enhancer). (A) *PAX6*-SIMO driving eGFP and *PAX6*-7CE3 driving mCherry. (B) Dye-swap experiment with *PAX6*-SIMO driving mCherry and *PAX6*-7CE3 driving eGFP. In both A and B, eGFP and mCherry expression is observed in both lens and hindbrain indicating complete crosstalk of activity between the two CREs consistent with the lack of insulation.

**Supplementary fig. 4:**
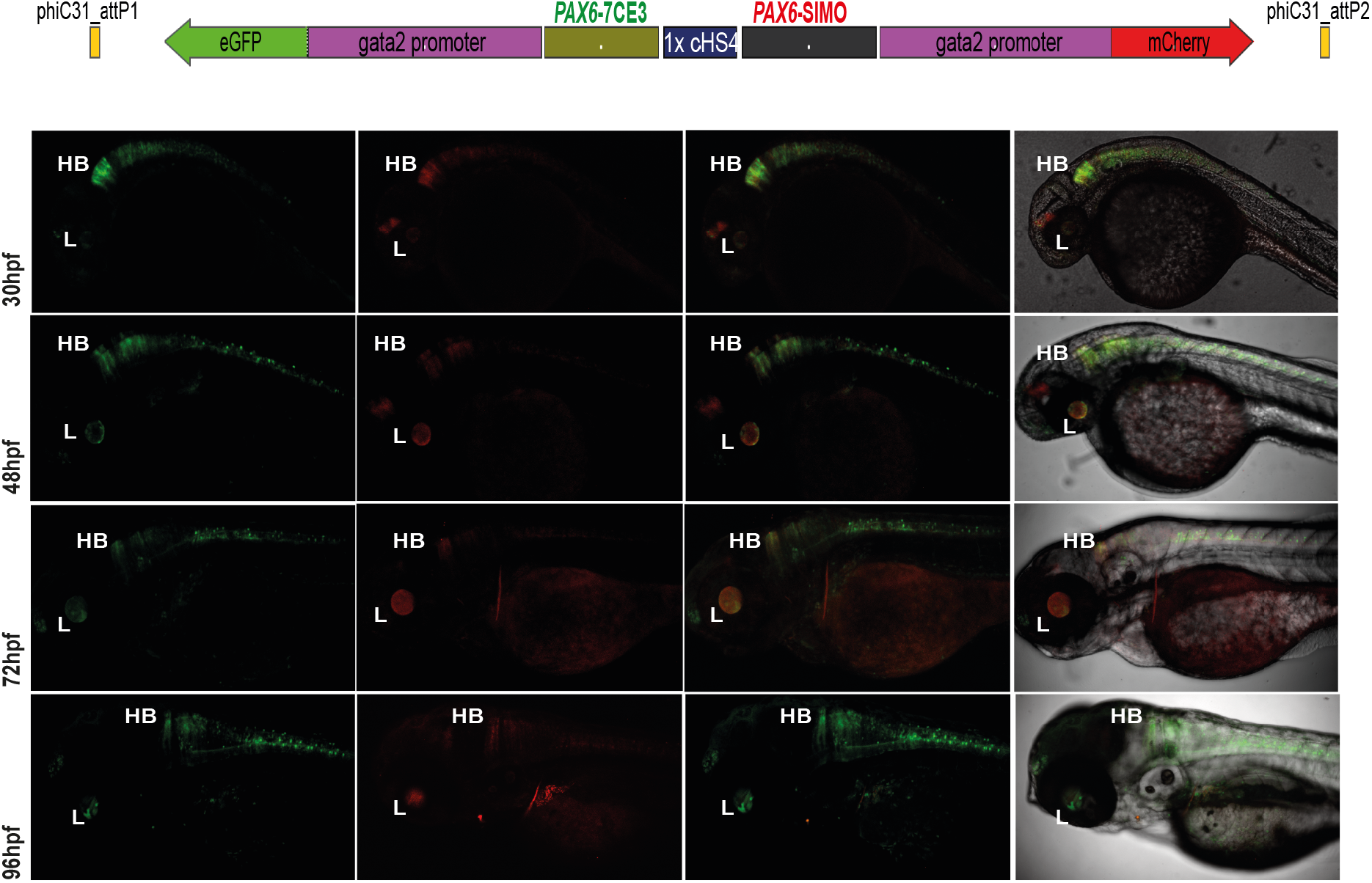
Assessment of tissue-specific CRE activity from the dual-CRE dual-reporter replacement construct with one copy of insulator sequence. Replacement constructs designed with previously well-characterised enhancers from *PAX6* locus (*PAX6*-7CE3, hindbrain enhancer and *PAX6*-SIMO, lens enhancer). Confocal images of F1 embryos (30-96hpf) derived from founder lines injected with the replacement cassettes bearing the enhancer-reporter cassettes separated by one copy of the insulator sequence (1XcHS4). eGFP and mCherry expression is observed in both lens (L) and hindbrain (HB) indicating a complete crosstalk of activity between the two CREs.

**Supplementary fig. 5:**
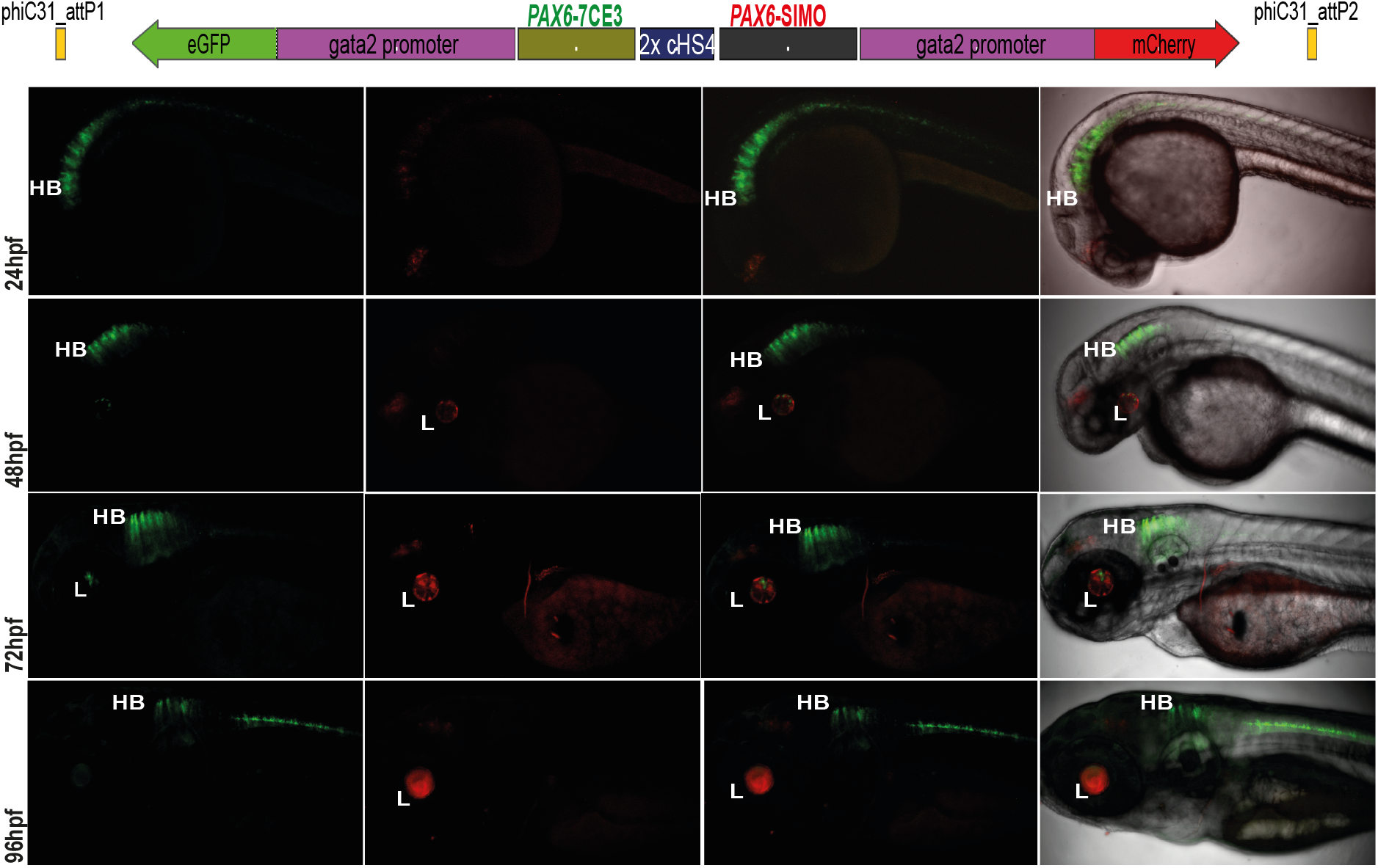
Assessment of tissue-specific CRE activity from the dual-CRE dual-reporter replacement construct with two copies of insulator sequence. Replacement constructs designed with previously well-characterised CREs from *PAX6* locus (*PAX6*-7CE3, hindbrain enhancer and *PAX6*-SIMO, lens enhancer). Confocal images of F1 embryos (24-96hpf) derived from founder lines injected with the replacement cassettes bearing the CRE-reporter cassettes separated by two copies of the insulator sequence (2XcHS4). eGFP and mCherry expression is observed largely restricted to either lens (L) or hindbrian (HB) indicating a blocking of crosstalk of activity between the two CREs by the presence of two copies of insulator sequence.

**Supplementary fig. 6:**
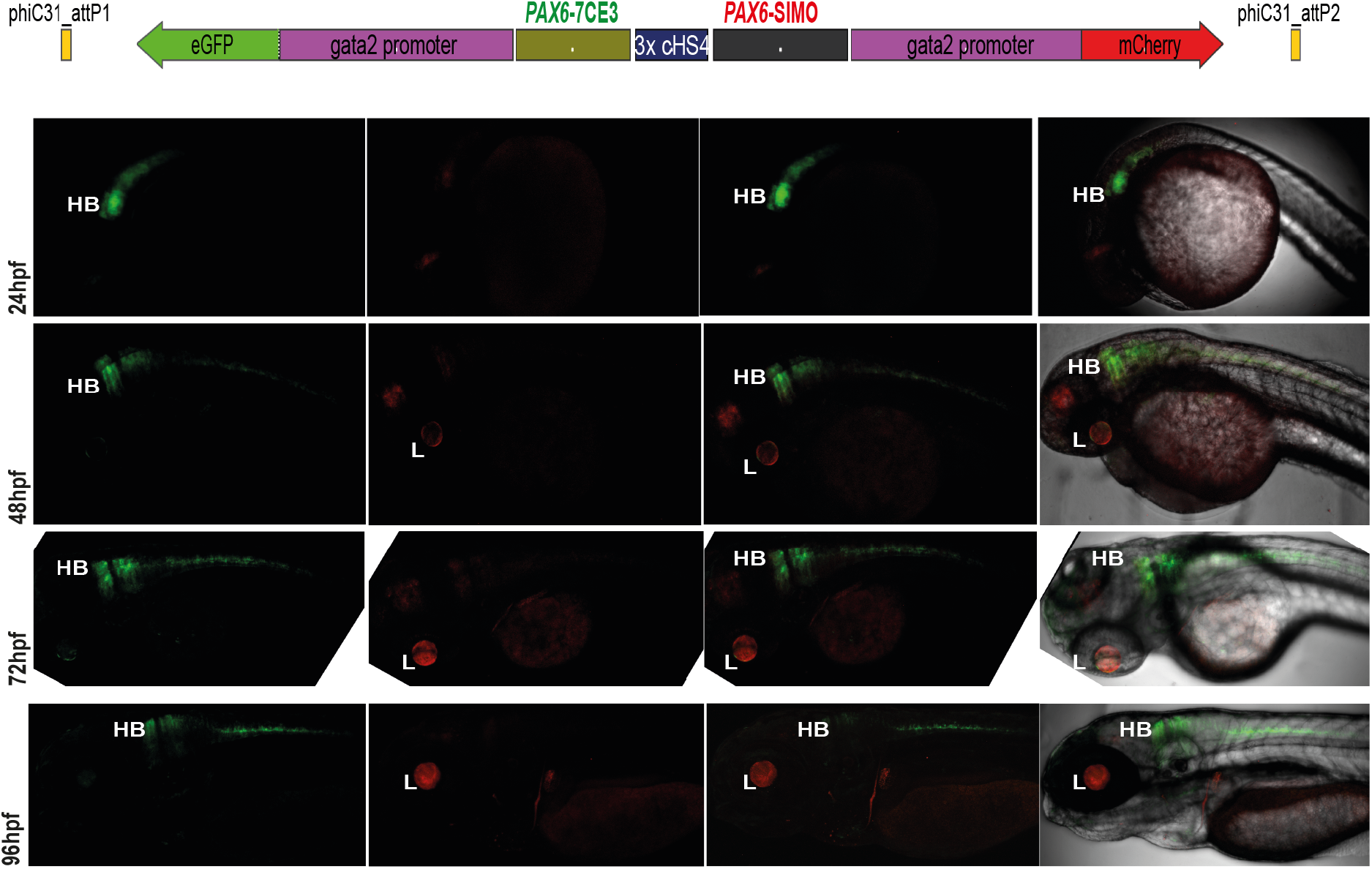
Assessment of tissue-specific CRE activity from the dual-CRE dual-reporter replacement construct with three copies of insulator sequence. Replacement constructs designed with previously well-characterised CREs from *PAX6* locus (*PAX6*-7CE3, hindbrain enhancer and *PAX6*-SIMO, lens enhancer). Confocal images of F1 embryos (24-96hpf) derived from founder lines injected with the replacement cassettes bearing the CRE-reporter cassettes separated by three copies of the insulator sequence (3XcHS4). eGFP and mCherry expression is observed completely restricted to either lens (L) or hindbrian (HB) indicating a complete insulation of crosstalk of activity between the two CREs by the presence of three copies of insulator sequence.

**Supplementary movie 1:** Confocal imaging of 30hpf embryo derived from transgenic line bearing the *Shh*-SBE2gfp/3XcHS4/*Shh*-SBE4mCherry replacement construct. The distinct expression domains of SBE2 and SBE4 enhancers in the developing forebrain are seen in green and red respectively, while the region where their activities overlap is depicted in yellow.

**Supplementary movie 2:** Time-lapse video of embryo derived from transgenic line bearing the *Shh*-SBE2gfp/3XcHS4/*Shh*-SBE4mCherry replacement construct. Images were acquired from 54hpf-69hpf, with a time interval of one hour. The distinct-expression domains of SBE2 and SBE4 enhancers in the developing forebrain are seen in green and red respectively.

**Supplementary movie 3:** Time-lapse video of embryo derived from transgenic line bearing the *SHH*-SBE2-Wtgfp/3XcHS4/*SHH*-SBE2-Mut-mCherry replacement construct. Images were acquired from 40hpf-60hpf, with a time interval of two hours.

**Supplementary table 1:** Details of oligonucleotides used in the study for generation of landing pads and replacement constructs, and mapping of site of integration of transgene in landing lines.

**Supplementary table 2:** Overview of transgenic lines generated in the study

**Supplementary table 3:** Quantification data of eGFP and mCherry intensities in transgenic lines bearing the replacement constructs described in the study.

